# Taxonomic Uncertainty and the Anomaly Zone: Phylogenomics Disentangle a Rapid Radiation to Resolve Contentious Species (*Gila robusta* complex) in the Colorado River

**DOI:** 10.1101/692509

**Authors:** Tyler K. Chafin, Marlis R. Douglas, Max R. Bangs, Bradley T. Martin, Steven M. Mussmann, Michael E. Douglas

## Abstract

Species is an indisputable unit for biodiversity conservation, yet their delimitation is fraught with both conceptual and methodological difficulties. A classic example is the taxonomic controversy surrounding the *Gila robusta* complex in the lower Colorado River of southwestern North America. Nominal species designations were originally defined according to weakly diagnostic morphological differences that conflicted with traditional genetic analyses. Consequently, the complex was re-defined as a single polytypic unit, with the proposed ‘threatened’ status of two being withdrawn at the federal level. Here, we utilized dense spatial and genomic sampling (N=387 and >22k loci) to re-evaluate the status of the complex, based on SNP-based coalescent and polymorphism-aware phylogenetic models. In doing so, all three species were supported as evolutionarily independent lineages, despite widespread phylogenetic discordance. To understand this discrepancy with past studies, we categorized evolutionary mechanisms driving discordance. We tested (and subsequently rejected) prior hypotheses suggesting that phylogenetic discord in the complex was hybridization-driven. Instead, we found the *G. robusta* complex to have diverged within the ‘anomaly zone’ of tree space and, as such, have accumulated inconsistent patterns of diversity which have confounded prior studies. After extending these analyses with phylogeographic modeling, we propose that this is reflective of a rapid radiation promoted by Plio-Pleistocene tectonism. Our results not only support resurrection of the three species as distinct entities, but also offer an empirical example of how phylogenetic discordance can be categorized in other recalcitrant taxa where variation is primarily partitioned at the species-level.

## Introduction

Complex evolutionary histories remain consistently difficult to disentangle, despite a recent paradigm shift towards the development of increasingly comprehensive datasets (e.g. Edwards 2009; Giarla and Esselstyn 2015). Regardless of these efforts, phylogenetic uncertainty is still prevalent, and with wide-ranging consequences on the study of macroevolutionary patterns (Stadler et al. 2016; Pereira and Schrago 2018), trait evolution (Hahn and Nakhleh 2016; Mendes et al. 2016; Wu et al. 2018), and ecological and biogeographic processes (Rangel et al. 2015; McVay et al. 2017).

Importantly, phylogenetic uncertainty also translates to taxonomic uncertainty. This is because modern systematic taxonomy fundamentally describes homology [i.e. Darwin’s (1859) ‘propinquity of descent’ (Simpson 1961)], which, by definition, requires a phylogenetic context. Phylogenetic uncertainty in this sense can manifest itself as a soft polytomy (= ‘honest’ uncertainty), the erroneous promotion of non-monophyletic clades, or controversial ‘splitting’ *versus* ‘lumping.’ Incomplete or biased sampling is often a driver of this disparity (Ahrens et al. 2016; Reddy et al. 2017). Here, narrow taxon sampling may introduce substantial ascertainment bias (=systematic deviations due to sampling). On the other hand, a broader yet sparse sampling regime often fails to sample cryptic lineages (Heath et al. 2008) — with subsequent impacts on both the delimitation of species (Pante et al. 2015; Linck et al. 2019) and study of their traits (Beaulieu and O’Meara 2018).

These sources of uncertainty culminate in topologies that often fluctuate with regard to sampling designs or methodologies, and this translates into taxonomic uncertainty [e.g. Ctenophora *versus* Porifera as sister to all other animals (Pisani et al. 2015; Whelan et al. 2015; Simion et al. 2017)]. Access to genome-scale data has alleviated some of these issues by offering a level of precision not possible with single-gene phylogenies (Philippe et al. 2005). However, their inherent complexity and heterogeneity introduces new problems, and consequently, additional sources of phylogenetic uncertainty.

Gene tree heterogeneity is an ubiquitous source of discordance in genomic data, and ‘noise’ as a source of this variance must consequently be partitioned from ‘signal’ (where ‘noise’ is broadly categorized as systematic or stochastic error). Large genomic datasets can reduce stochastic error (Kumar et al. 2012), yet it still remains a prevalent issue when individual genes are examined (Springer and Gatesy 2016). On the other hand, systematic error in phylogenomics may represent a probabilistic bias towards incongruence that is inherent to the evolutionary process itself (Maddison 1997). This, in turn, exemplifies the complications introduced by genomic data: As genomic resolution increases, so also does the probability of sampling unmodeled processes (Rannala and Yang 2008; Lemmon and Lemmon 2013). This potential (i.e., simultaneously decreasing stochastic error as systematic error increases) produces the very real possibility of building a highly supported tree that is ultimately incorrect.

Certain demographic histories are more predisposed to systematic error than others. For instance, when effective population sizes are large and speciation events exceptionally rapid, time between divergence events may be insufficient to sort ancestral variation, such that the most probable gene topology will conflict with the underlying species branching pattern. This results in what has been coined an ‘anomaly zone’ of tree space (i.e., dominated by anomalous gene trees (AGTs); (Degnan and Rosenberg 2006). Inferring species trees is demonstrably difficult in this region (Liu and Edwards 2009), and exceedingly so if additional sources of phylogenetic discordance, such as hybridization, are also apparent (Bangs et al. 2018). Here, historical or persistent gene flow both compresses apparent divergence in species-trees (Leache et al. 2014b) and similarly drives a predominance of AGTs which can supercede ‘correct’ branching patterns in some regions of parameter space (Long and Kubatko 2018). The result is a confounding effect on the adequate delineation of phylogenetic groupings (e.g., a necessary step of biodiversity conservation), as well as a limitation in the downstream analysis of affected species trees (Bastide et al. 2018; Luo et al. 2018; Morales and Carstens 2018; Bangs et al. 2020).

In clades with such complex histories, it is often unclear where the source of poor support and/or topological conflict resides (Richards et al. 2018). To analytically account for gene tree conflict, it is necessary to categorize these sources and select approaches accordingly. Failure to do so promotes a false confidence in an erroneous topology, as driven by model misspecification (Philippe et al. 2011). The overwhelmingly parametric nature of modern phylogenetics ensures that imperative issues will revolve around the processes being modeled, and what they actually allow us to ask from our data (Sullivan and Joyce 2005). However, the selection of methods that model processes of interest requires an *a priori* hypothesis that delimits which processes are involved. Diagnosing prominent processes is difficult in that a phylogenetic context is required from which to build such hypotheses. Fortunately, a wealth of information can be parsed from otherwise ‘non-phylogenetic’ signal (*sensu* Philippe et al. 2005). For example, many statistical tests diagnose hybridization via its characteristic signature on the distribution of discordant topologies (e.g. Pease and Hahn 2015). Theoretical predictions regarding AGTs and the parameters under which they are generated are also well characterized (Degnan and Salter 2005; Degnan and Rosenberg 2009). Thus, by applying appropriate analytical approaches that sample many independently segregating regions of the genome, empiricists can still derive biologically meaningful phylogenies, despite the presence of complicated species-histories (McCormack et al. 2009; Kumar et al. 2012).

Here, we demonstrate an empirical approach that infers species-histories and sources of subtree discordance when conflict originates not only from anomaly zone divergences but also hybridization. To do so, we used SNP-based coalescent and polymorphism-aware phylogenetic methods (Chifman and Kubatko 2014; Leache et al. 2014a; De Maio et al. 2015) that bypass the necessity of fully-resolved gene trees. We combine coalescent predictions, phylogenetic network inference (Solís-Lemus and Ané 2016), and novel coalescent phylogeographic methods (Oaks 2018) to diagnose the sources of phylogenetic discordance and, by so doing, resolve a seemingly convoluted complex of study-species (the *Gila robusta* complex of the lower Colorado River basin). We then contextualize our results to demonstrate the downstream implications of ‘problematic’ tree-space for threatened and endangered taxa, as represented by our study complex.

### Gila

Few freshwater taxa have proven as problematic in recent years as the *Gila robusta* complex (Cyprinoidea: Leuciscidae) endemic to the Gila River basin of southwestern North America (Fig. 1). The taxonomic debate surrounding this complex exemplifies an inherent conflict between the traditional rigidity of systematic taxonomy *versus* the urgency of decision-making for conservation and management (Forest et al. 2015). Our study system is the Gila River, a primary tributary of the lower basin Colorado River that drains the majority of Arizona and ∼11% of New Mexico. The critical shortage of water in this region (Sabo et al. 2010) is a major geopolitical driver for the taxonomic controversy surrounding the study species. As an example, the lower Colorado River basin supplies approximately half of the total municipal and agricultural water requirements of the state of Arizona, and nearly two-thirds of its total gross state product (GSP) (Bureau of Reclamation 2012; James et al. 2014). This disproportionate regional reliance creates tension between the governance of a resource and its usage (e.g. Huckleberry and Potts 2019) which in turn magnifies the stakes involved in conservation policy (Minckley 1979; Carlson and Muth 1989; Minckley et al. 2006).

**Figure 1:**
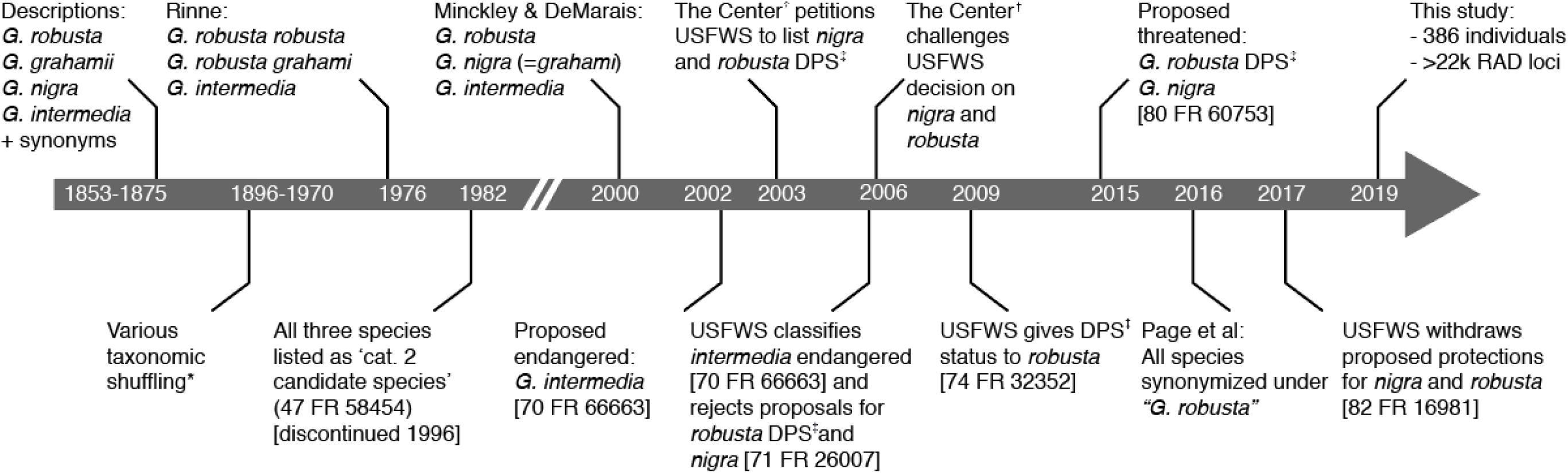
Timeline of the conservation status of *Gila* species endemic to the lower Colorado River basin [*See Copus et al (2018) for a detailed overview of taxonomic synonymies; †’The Center’ refers to the Center for Biological Diversity (501c3), Tuscon, AZ; ‡’DPS’ = Distinct Population Segment as referenced in the Endangered Species Act (ESA 1973; 16 U.S.C. § 1531 et seq), here referring specifically to a lower basin sub-unit of *G. robusta*]. Note that timeline is not to scale.

We focused on three species (Roundtail chub, *G. robusta*; Gila chub, *G. intermedia*; and Headwater chub, *G. nigra*) that comprise a substantial proportion of the endemic ichthyfauna of the Gila Basin [=20% of 15 extant native species (excluding extirpated *G. elegans*, *Ptychocheilus lucius*, and *Xyrauchen texanus*); Minckley and Marsh 2009]. Historically, the focal taxa have been subjected to numerous taxonomic rearrangements (Fig. 1). Until recently, the consensus was defined by Minckley and DeMarais (2000) on the basis of morphometric and meristic characters. These have since proven of limited diagnostic capacity in the field, thus provoking numerous attempts to re-define morphological delimitations (Brandenburg et al. 2015; Moran et al. 2017; Carter et al. 2018). Genetic evaluations have to date been inconclusive (Schwemm 2006; Copus et al. 2018), leading to a recent taxonomic recommendation that subsequently collapsed the complex into a single polytypic species (Page et al. 2016, 2017).

## Methods

### Taxonomic Sampling

A representative panel of *N*=386 individuals (Table S1; Fig. 2) was chosen from existing collections (Douglas et al. 2001; Douglas and Douglas 2007), to include broad geographic sampling of the complex as well as congeners. For the sake of clarity, we employ herein the nomenclature of Minckley and DeMarais (2000)., and retained species-level nomenclature for all members of the *Gila robusta* complex. Additionally, we discriminate between *G. robusta* from the upper and lower basins of the Colorado River ecosystem (Chafin et al. 2019)

**Figure 2:**
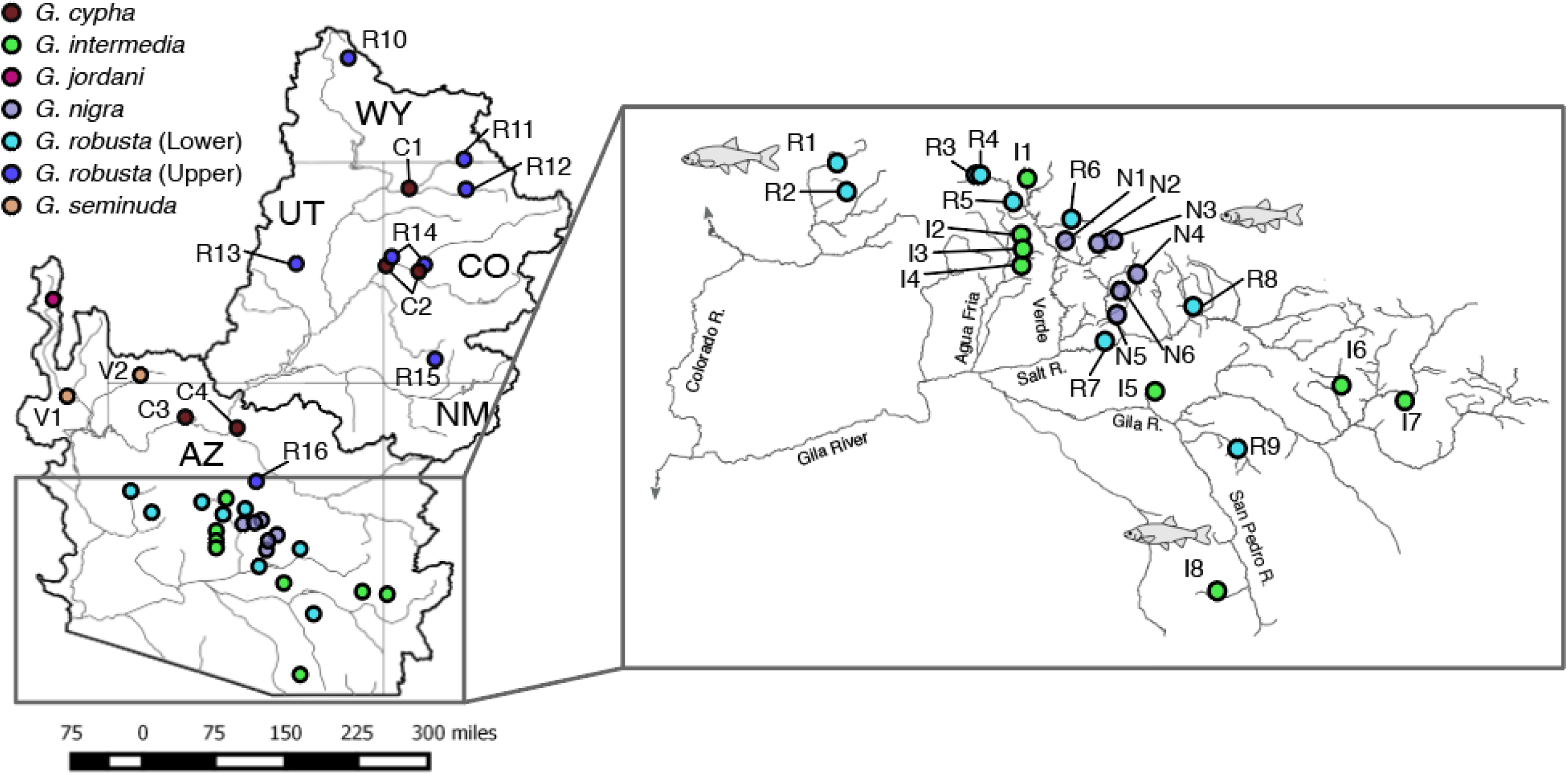
Sampling localities for *Gila* within the Colorado River Basin, southwestern North America. Locality codes are defined in Table S1. Sympatric locations (R14 and C2) are slightly offset for visibility purposes. Map insert increases the viewing scale for sampling sites within the lower basin ‘complex’ (Bill Williams and Gila rivers).

No self-sustaining populations of wild *Gila elegans* exist, thus samples were provided by the Southwestern Native Aquatic Resources and Recovery Center (Dexter, NM). The genus *Ptychocheilus* served to root the *Gila* clade within the broader context of western leuciscids (Schönhuth et al. 2012, 2014, 2018).

### Reduced-Representation Sequencing

Genomic DNA was extracted using either PureGene® or DNeasy® kits (Qiagen Inc.) and quantified via fluorometer (Qubit™; Thermo-Fisher Scientific). Library preparations followed the published ddRAD protocol (Peterson et al. 2012). Restriction enzyme and size-selection ranges were first screened using an *in silico* procedure (Chafin et al. 2018), with the target fragment sizes further optimized by quantifying digests for 15 representative samples on an Agilent 2200 TapeStation. Final library preparations were double-digested using a high-fidelity *PstI* (5’-CTGCAG-3’) and *MspI* (5’-CCGG-3’) following manufacturer’s protocols (New England Biosciences). Digests were purified using bead purification (Ampure XP; Beckman-Coulter Inc.), and standardized at 100 ng per sample. Samples were ligated with customized adapters containing unique in-line barcodes, pooled in sets of 48, and size-selected at 250-350bp (not including adapter length), using a Pippin Prep automated gel extraction instrument (Sage Sciences). Adapters were then extended in a 12-cycle PCR using Phusion high-fidelity DNA polymerase (New England Biosciences Inc.), completing adapters for Illumina sequencing and adding an i7 index. Libraries were pooled to N=96 samples per lane (i.e., 2 sets of 48) for 100bp single-end sequencing on an Illumina HiSeq 2500 at the University of Wisconsin Biotechnology Center (Madison, WI).

### Data Processing and Assembly

Raw Illumina reads were demultiplexed and filtered using the pyRAD pipeline (Eaton 2014). We removed reads containing >1 mismatch in the barcode sequence, or >5 low-quality base-calls (Phred Q<20). Assembly of putative homologs was performed using *de novo* clustering in VSEARCH (Rognes et al. 2016) using an 80% mismatch threshold. Loci were excluded according to following criteria: >5 ambiguous nucleotides; >10 heterozygous sites in the alignment; >2 haplotypes per individual; <20X and >500X sequencing depth per individual; >70% heterozygosity per-site among individuals.

Our ddRAD approach generated 22,768 loci containing a total of 173,719 variable sites, of which 21,717 were sampled (=1/ locus). Mean per-individual depth of coverage across all retained loci was 79X. All relevant scripts for post-assembly filtering and data conversion are available as open-source (github.com/tkchafin/scripts).

### Phylogenetic Inference

We formulated two simple hypotheses with regards to independent evolutionary sub-units. If populations represented a single polytypic species, then phylogenetic clustering should reflect intraspecific processes (e.g. structured according to stream heirarchy; Meffe and Vrijenhoek 1988). However, if *a priori* taxon assignments are evolutionarily independent, then they should be recapitulated in the phylogeny. Given well-known issues associated with application of supermatrix/ concatenation approaches (Degnan and Rosenberg 2006; Edwards et al. 2016) and pervasive gene-tree uncertainty associated with short loci (Leaché and Oaks 2017), we also employed SNP-based methods that bypassed the derivation of gene trees (Leaché and Oaks 2017).

We first explored population trees in SVDquartets (Chifman and Kubatko 2014, 2015; as implemented in PAUP*, Swofford 2002) across 12 variably filtered datasets using four differing occupancy thresholds per SNP locus (i.e., 10, 25, 50, and 75%), along with three differing thresholds per individual (10, 25, and 50%). These filtered datasets ranged from 7,357– 21,007 SNPs, with 8.48–43.65% missing data and 256–347 individuals. SVDquartets eases computation by inferring coalescent trees from randomly sampled quartets of species (i.e. optimizing among 3 possible unrooted topologies). It then generates a population tree with conflicts among quartet trees minimized via implementation of a quartet-assembly algorithm (Snir and Rao 2012). Given run-time constraints (the longest was 180 days on 44 cores), all runs sampled 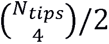 quartets and were evaluated across 100 bootstrap pseudo-replicates.

We also used a polymorphism-aware method (PoMo; Schrempf et al. 2016) in IQ-Tree (Nguyen et al. 2014). PoMo considers allele frequencies rather than single nucleotides, thus allowing evaluation of change due to both substitution and drift. To provide PoMo with empirical estimates of polymorphism, we used the entire alignment, to include non-variable sequences. We filtered liberally using individual occupancy thresholds of 10% per-locus so as to maximize individual retention and per-population sample sizes. We then deleted populations that contained <2 individuals, and loci with >=90% missing data per-population. This yielded a dataset of 281,613 nucleotides and 40 tips. Non-focal outgroups were excluded due to their disproportionate effect on missing data. Analyses were pseudo-replicated across 100 bootstraps.

We also calculated concordance factors (CFs) using a Bayesian concordance analysis in BUCKy (Larget et al. 2010), parallelized across all quartets via an adaptation of the TICR pipeline (Stenz et al. 2015). To prepare these data, we sampled all non-monomorphic full gene alignments for which at least 1 diploid genotype could be sampled per population. We excluded outgroups and non-focal *Gila* so as to maximize number of loci retained. This yielded 3,449 genes across 31 sampled tips. Gene-tree priors were generated using MrBayes v.3.2.6 (Ronquist et al. 2012) with 4 independent chains, each of which was sampled every 10,000 iterations, with a total chain length of 100,000,000 iterations and 50% discarded as burn-in. BUCKy was then run in parallel to generate quartet CFs across 31,465 quartets, using a chain length of 10,000,000, again with 50% burn-in. Quartet topologies were used to generate a population tree using QuartetMaxCut (Snir and Rao 2012), using the *get-pop-tree.pl* script from TICR (Stenz et al. 2015; https://github.com/nstenz/TICR).

### Comparing Phylogenies and Estimating Site-wise Conflict

To evaluate the performance of SVDquartets, TICR, and PoMo, we first computed site-wise log-likelihood scores (*SLS*) for each topology by performing a constrained ML search in IQ-Tree. For comparison, we also generated an unconstrained concatenated tree. All ML analyses employed a GTR model with empirical base frequencies and gamma-distributed rates, and were assessed across 1,000 bootstrap pseudoreplicates. Analyses were also reduced to a subset of tips common across all variably filtered datasets. We quantified the phylogenetic signal supporting each resolution as the difference in site-wise log-likelihood scores (Δ*SLS*) between each population tree and the concatenation tree (Shen et al. 2017). We then calculated site-wise concordance factors (s*CF*) as an additional support metric (Minh et al. 2018).

### Tests of Hybridization and Deep-Time Reticulation

*D*-statistics (Green et al. 2010; Eaton and Ree 2013) were calculated using Comp-D (Mussmann et al. 2020). To further test hypotheses of reticulation, we used quartet CFs as input for phylogenetic network inference using the SNaQ algorithm in PhyloNetworks (Solís-Lemus and Ané 2016; Solís-Lemus et al. 2017). The network was estimated under models of 0−5 hybrid nodes (*h*). Models were evaluated using 48 independent replicates, with the best-fit model being that which maximized change in pseudolikelihood. Given the computational constraints of network inference, we reduced the dataset to N=2 populations per focal species (=12 total tips). We also explicitly tested for putative hybrid taxa using HyDe (Blischak et al. 2018), which rapidly tests all possible parent-descendant combinations to diagnose hybrid lineages using phylogenetic invariants. In this case, hybrids are detected by considering the ratio of two phylogenetic invariants which evaluate to zero for opposing topologies (Meng and Kubatko 2009; Chifman and Kubatko 2015; Kubatko and Chifman 2016). This ratio is incorporated into what Kubatko & Chifman (2016) refer to as the Hils statistic, *H*, which is compared with a normal distribution for hypothesis testing in hybrid taxa. Significance was assessed at a Bonferroni-corrected threshold = 5.7 × 10^−8^. Finally, we generated distance-based networks using the NeighborNet algorithm (Bryant and Moulton 2004), as implemented by SplitsTree4 (Huson 1998).

### Anomaly Zone Detection

Coalescent theory characterizes the boundaries of the anomaly zone in terms of branch lengths in coalescent units (Degnan and Rosenberg 2006). To test if contentious relationships in our tree fell within the anomaly zone, we first transformed branch lengths using quartet CFs (Stenz et al. 2015, equation 1), then tested if internode branch lengths fell within the theoretical boundary for the anomaly zone (Linkem et al. 2016, equation 1). Code for these calculations are modified from Linkem et al. (2016) and are available as open-source (github.com/tkchafin/anomaly_zone).

### Tests of Co-divergence

The contemporary course of the Colorado River resulted from the Pliocene erosion of the Grand Canyon and subsequent connection of the modern-day upper and lower basins, to include stream capture of the Gila River (McKee et al. 1967; Minckley et al. 1986). *Gila* in the lower Colorado River basin then differentiated following one or more colonization events (e.g. Rinne 1976). Subsequent work (Douglas et al. 1999) supported this conclusion by examining contemporary phenotypic variation among all three species as a function of historical drainage connectivity, with the conclusion that body shape was most readily explained by Pliocene hydrography.

We tested if divergences were best explained by a model of *in situ* diversification following a single colonization event, or instead by multiple, successive colonizations. We compared divergence models using a Bayesian approach (program Ecoevolity, Oaks 2018) that used a coalescent model (Bryant et al. 2012) to update a prior expectation for the number of evolutionary events across independent comparisons. Four independent MCMC chains were run with recommended settings and a burn-in that maximized effective sample sizes. Event models followed a Dirichlet process, with the concentration parameter exploring four alternative gamma distributed priors (i.e. *α*=2.0, *β*=5.70; *α*=0.5, *β*=8.7; *α*=1.0, *β*=0.45; and *α*=2.0, *β*=2.18).

We randomly sampled 2,000 full-locus alignments, then examined potential co-divergences in the lower-basin complex by selecting a series of pairwise comparisons: *Gila elegans* x *G. robusta* (lower); *G. seminuda* x *G. robusta* (lower); *G. jordani* x *G. robusta* (lower); *G. intermedia* x *G. robusta* (lower); and *G. intermedia* x *G. nigra* (lower). These targeted nodes represent F, G, H, I, and N in the SVDQuartets topology (Fig. 3A).

**Figure 3:**
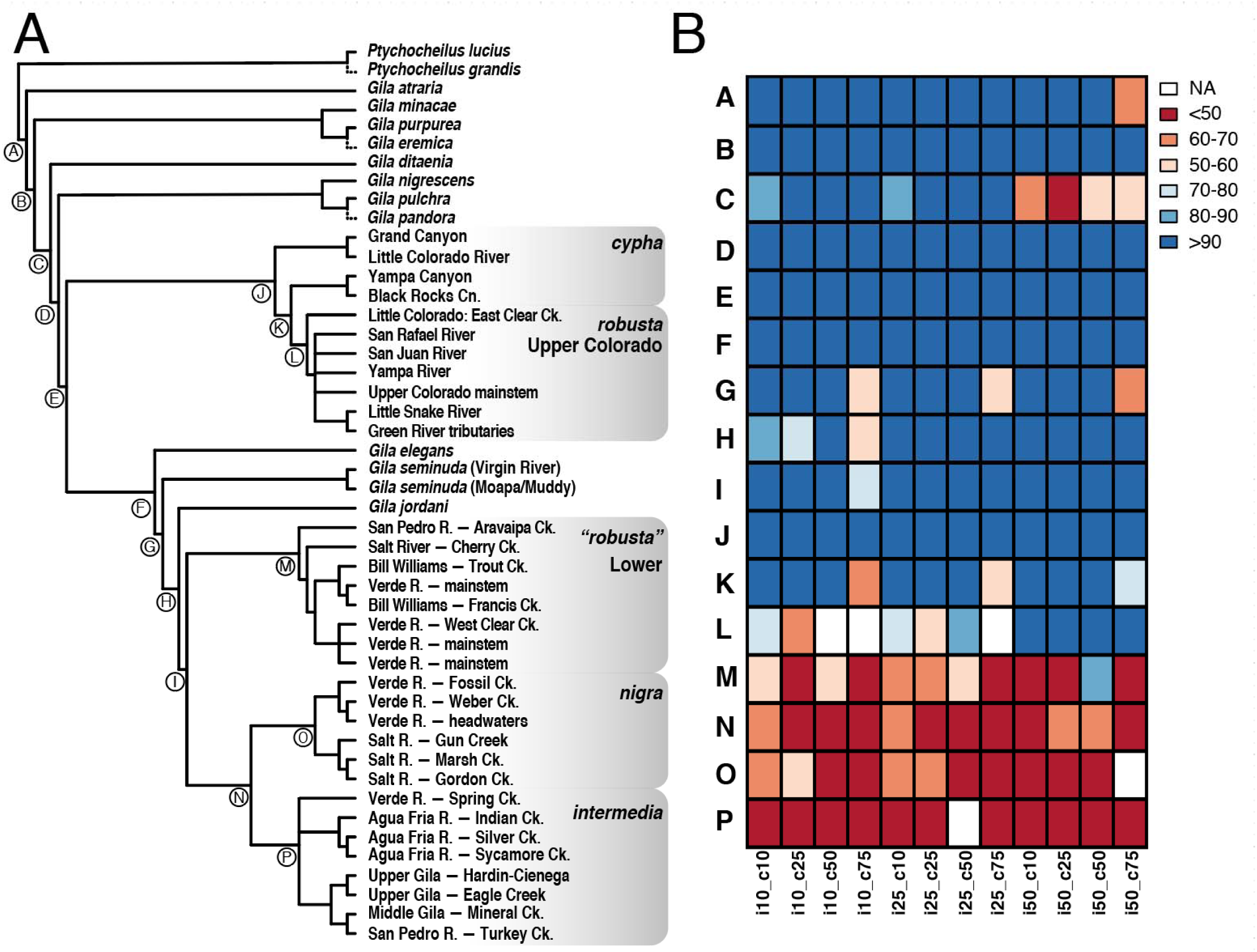
(A) Majority-rule consensus cladogram of SVDQuartets across 12 variably filtered SNP datasets varying from 7,357–21,007 SNPs and 256–347 individuals. (B) Binned bootstrap concordance values are reported for each dataset, coded by the matrix occupancy threshold per individual (“i”) and per column (“c”; e.g. i50_c50 = 50% occupancy required per individual and per column). Dashed terminal branches indicate positions for taxa missing from >50% of datasets. For detailed locality information, refer to Table S1.

## Results

### Phylogenetic Conflict in Gila

Tree reconstructions across all three population methods were relatively congruent (SVDquartets = Fig. 3; PoMo, and TICR = Fig. 4). The concatenated supermatrix tree (Fig. S1) was also largely congruent with the population trees, but with two major disparities (discussed below). Bootstrap support was variable and declined with decreasing nodal depth in the SVDquartets analysis (Fig. 3), whereas the vast majority of nodes in PoMo were supported at 100% (Fig. 4A).

**Figure 4:**
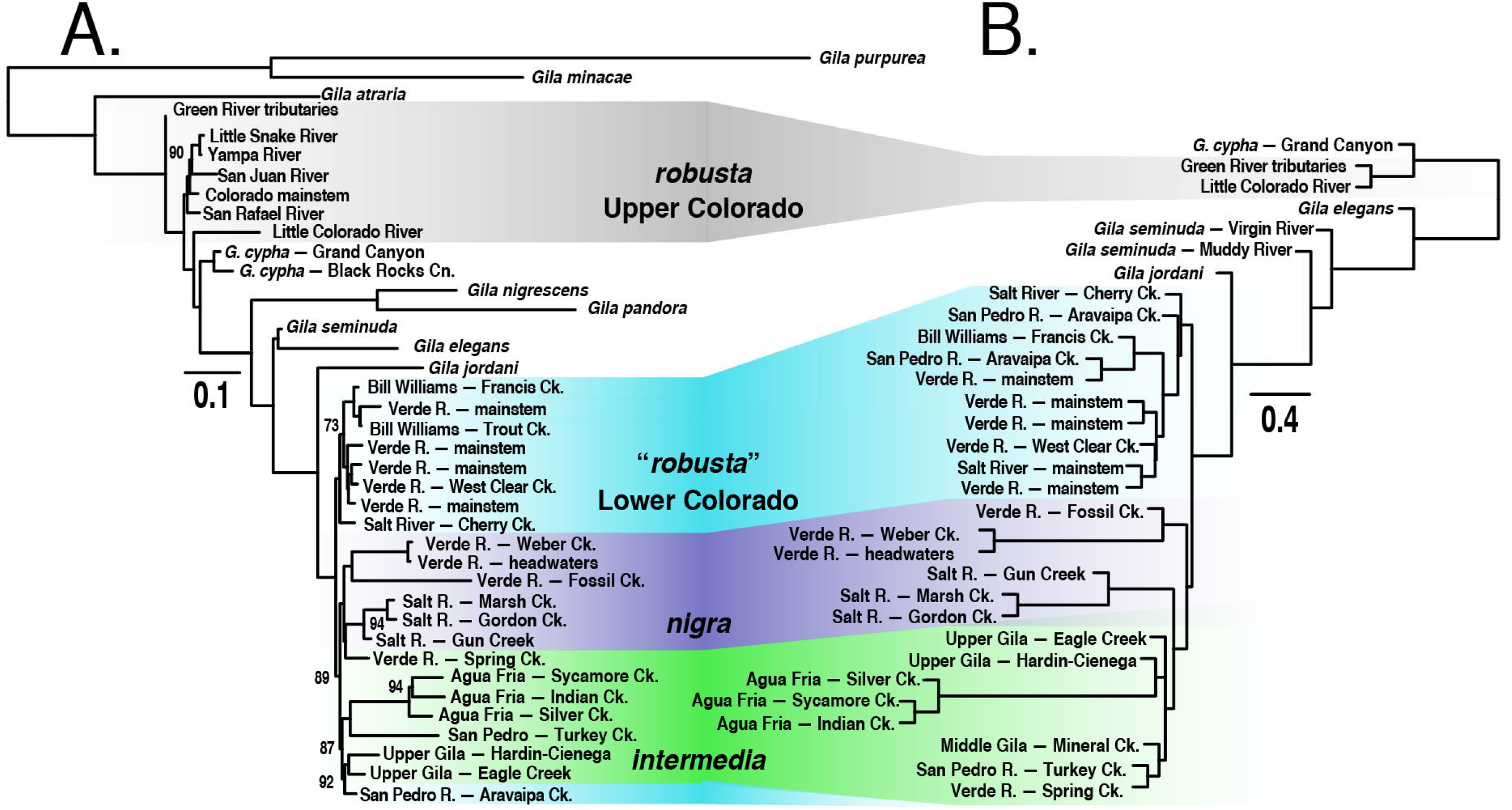
(A) PoMo phylogram with branch lengths as the number of substitutions *and* inferred number of drift events per site, with branch supports (as values <100%) representing concordance among 1,000 bootstrap replicates, inferred using a dataset consisting of 281,613 nucleotides and 40 tips; (B) TICR phylogram reporting branch lengths in coalescent units, calculated from 31,465 quartets evaluated across 3,449 full alignments of ddRAD loci. For detailed locality information, refer to Table S1.

All analyses consistently supported the monophyly of a clade consisting of *G. intermedia*, *G. nigra*, and lower basin *G. robusta* (hereafter the ‘lower basin complex’). This clade had high bootstrap support in both SVDquartets and PoMo, and was universally placed as sister to *G. jordani*. *Gila robusta* was unequivocally polyphyletic in all analyses, forming two distinct groups geographically demarcated by the Grand Canyon. The lower basin *G. robusta* clade was monophyletic in all cases, save the concatenated tree, where it was paraphyletic (Fig. S1). It was also consistently recovered as sister to a monophyletic *G. nigra* + *G. intermedia*, with the exclusion of a single sample site (Aravaipa Creek) that nested within *G. intermedia* in the PoMo tree. Of note, this population had been previously diagnosed as trending towards *G. intermedia* in terms of morphology (Rinne 1976; DeMarais 1986), although hybridization was not supported by *D*-statistics (Table 1).

**Table 1:** Four-taxon *D*-statistic Tests of Admixture. Tests were performed for quartets sampled from *N*=386 *Gila* individuals. Results are reported across *N* separate quartet samples per four-taxon test, randomly sampled without replacement, with site patterns calculated from 21,717 unlinked SNPs. Significance is reported as the proportion of tests at *p*<0.05 (nSig/*N*) using chi-squared (*χ*^2^), *Z*-test^1^, and *Z*-test with Bonferroni correction^2^. Positive and negative values of *D* suggest introgression of the P3 lineage with either P2 or P1, respectively. Results in bold were also supported by the phylogenetic network. See Table S1 for detailed locality information.

| P3 | P2 | P1 | Mean $D$ | $N$ | nSig/$N$<br>($\chi^2$) | nSig/$N$<br>($Z^1$) | nSig/$N$<br>($Z^2$) |
| --- | --- | --- | --- | --- | --- | --- | --- |
| *cypha* | *jordani* | lower basin | 0.175 | 86400 | 0.033 | 0.072 | 0.001 |
| *cypha* | *seminuda* | lower basin | 0.099 | 86400 | 0.102 | 0.130 | 0.002 |
| *elegans* | *jordani* | lower basin | -0.063 | 84800 | 0.029 | 0.050 | 0.000 |
| *elegans* | *robusta* (lower) | *nigra/int.* | -0.026 | 413600 | 0.014 | 0.047 | 0.001 |
| *elegans* | *robusta* (upper) | *cypha* | -0.236 | 45056 | 0.380 | 0.415 | 0.045 |
| ***elegans*** | ***seminuda*** | lower basin | **0.302** | **86400** | **0.654** | **0.674** | **0.251** |
| *jordani* | *robusta* (lower) | *nigra/int.* | 0.087 | 601600 | 0.042 | 0.072 | 0.001 |
| *jordani* | *robusta* (lower) | *nigra/int.* | 0.091 | 212800 | 0.041 | 0.067 | 0.005 |
| *nigra* | *int.* (Salt) | *int.* (Verde) | 0.086 | 126976 | 0.057 | 0.082 | 0.001 |
| *robusta* (lower) | *intermedia* | *nigra* | 0.041 | 793600 | 0.001 | 0.002 | 0.000 |
| *robusta* (upper) | *jordani* | *robusta* (lower) | 0.165 | 168000 | 0.050 | 0.081 | 0.001 |
| *robusta* (upper) | *robusta* (lower) | *nigra/int.* | -0.009 | 601600 | 0.011 | 0.031 | 0.000 |
| *robusta* (upper) | *seminuda* | lower basin | -0.017 | 180800 | 0.030 | 0.053 | 0.004 |
| *seminuda* | *jordani* | lower basin | -0.204 | 81920 | 0.107 | 0.152 | 0.000 |
| *seminuda* | *robusta* (lower) | *nigra/int.* | 0.054 | 212800 | 0.011 | 0.031 | 0.001 |
| *atraria* | *robusta* (upper) | *cypha* | 0.082 | 57344 | 0.064 | 0.095 | 0.033 |
| *nigrescens* | *robusta* (lower) | *nigra/int.* | -0.075 | 485472 | 0.023 | 0.079 | 0.002 |
| *nigrescens* | *robusta* (upper) | *cypha* | -0.039 | 53248 | 0.040 | 0.066 | 0.005 |
| *pandora* | *robusta* (lower) | *nigra/int.* | -0.123 | 225600 | 0.012 | 0.105 | 0.010 |
| *pandora* | *robusta* (upper) | *cypha* | -0.047 | 24576 | 0.031 | 0.057 | 0.003 |

Topology within the *G. nigra* + *G. intermedia* clade was less consistent. Both were reciprocally monophyletic in the SVDquartets tree (albeit with low support; Fig. 3), whereas PoMo yielded a monophyletic *G. intermedia*, with but one population (Spring Creek) contained within *G. nigra* (Fig. 4A). The PoMo tree also conflicted with the other methods in its paraphyletic placement of upper basin *G. robusta*. We suspect this represents an artefact of well-known hybridization with sympatric *G. cypha* (Dowling and DeMarais 1993; Gerber et al. 2001; Douglas and Douglas 2007; Chafin et al. 2019).

### Discriminating Among Sources of Phylogenetic Conflict

Phylogenetic conflict was variably attributable to either hybridization or rapid divergence. We found support for a single reticulation event connecting *G. seminuda* and *G. elegans*, an hypothesis consistent with prior interpretations (DeMarais et al. 1992). This particular model (i.e., *h*=1) was selected as the one that maximized both the first [L’(*h*) = L(*h*) – L(*h*-1)] and second order [L’’(*h*) = L’(*h*+1) – L’(*h*)] rate of change in pseudolikelihood (Fig. S2-S3; following Evanno et al. 2005). Of note, introgression between *G. elegans* and *G. seminuda* was supported by elevated values of *h,* and by *D*-statistics (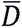 = 0.302 across 86,400 tests; Table 1). Likewise, distance-based networks (Fig. S3) supported this reticulation, as did *H*-tests from HyDe, where p-values supporting a hybrid origin for *G. seminuda* from *G. elegans* and lower-basin progenitors ranged from *p*=7.8×10^−9^ to *p*=5.6×10^−8^. Introgression between upper basin *G. robusta* and *G. cypha* was also supported (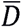 = −0.236 across 45,056 tests). No other introgressions were noted, thus rejecting the hypothesized hybrid origins for both *G. jordani* (Dowling and DeMarais 1993; Dowling and Secor 1997) and *G. nigra* (Demarais 1986; Minckley and DeMarais 2000).

Multiple internode pairs were observed in the anomaly zone (Fig. 5). In all cases, internode branches separating *G. nigra* and *G. intermedia,* and those separating their constituent lineages, reflected coalescent lengths that would yield anomalous gene trees. Not surprisingly, the internode separating *G. jordani* from the lower basin complex, and that of *G. robusta* from *G. intermedia*/ *G. nigra* (Fig. 5C; tan branches) also fell within the anomaly zone, per TICR and concatenated topology results.

**Figure 5:**
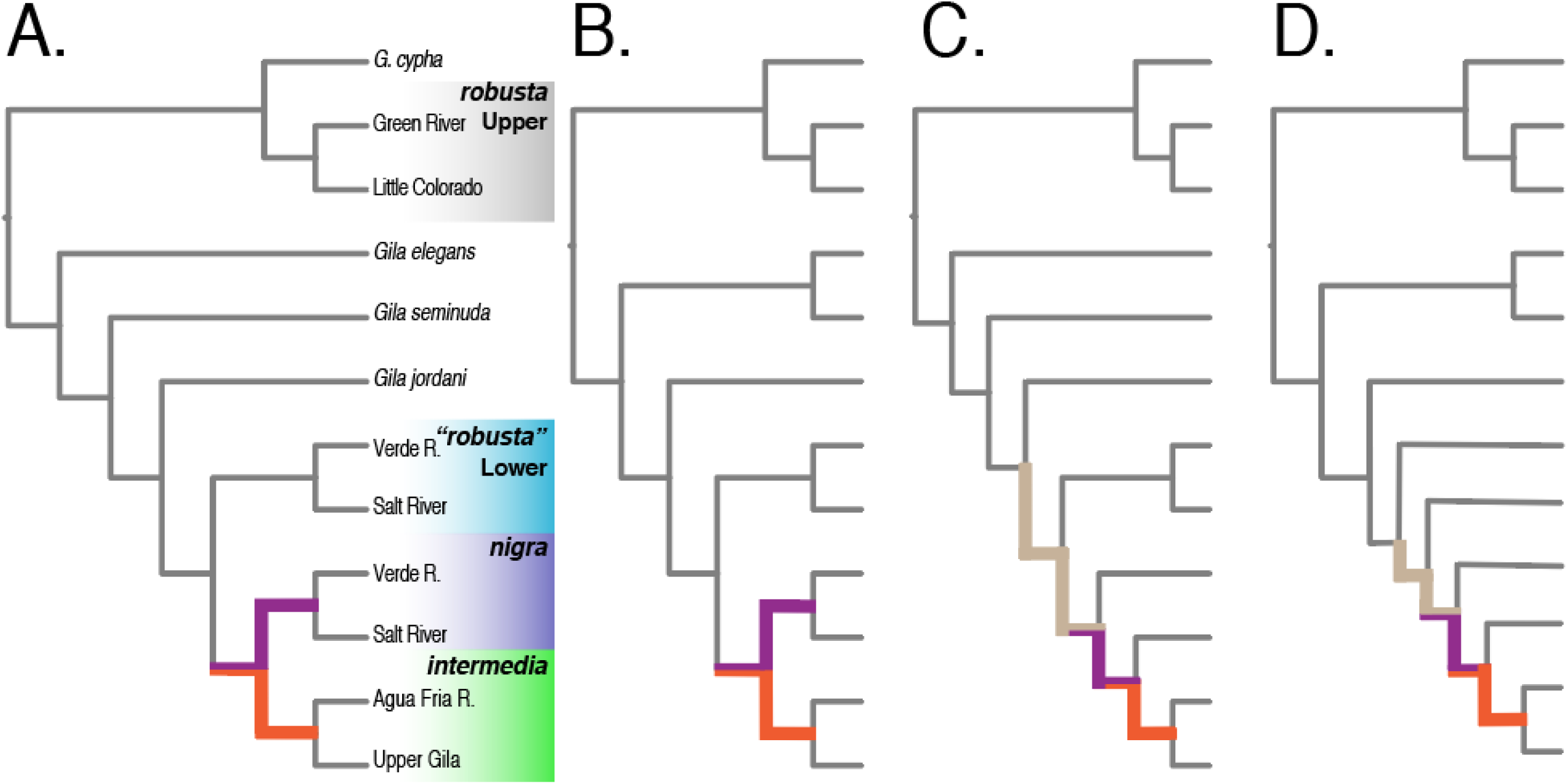
Internode pairs within the anomaly zone, as determined using coalescent-unit transformed branch lengths mapped onto the (A) SVDQuartets, (B) PoMo, (C) TICR, and (D) concatenated trees (displayed here as cladograms). Paired internodes are color-coded, with those bicolored indicating multiple anomalous divergences. For more detailed representations, refer to Figs. 3 and 4.

### Relative Performance of Species-Tree Methods

Change in site-likelihoods among constrained and unconstrained IQ-Tree searches in all cases suggested that the recovered species-trees were supported by a minority of sites (Fig. S4), an observation consistent with tree regions being in the anomaly zone. Several discrepancies also reflected idiosyncrasies among the different approaches. For example, the PoMo topology has a paraphyletic upper basin *G. robusta* within which *G. elegans*, *G. cypha*, *G. seminuda*, *G. jordani*, and the lower basin complex were subsumed (Fig. 4A). However, only ∼10% of SNPs supported this resolution (Fig. S5), a value far below the theoretical minimum s*CF* derived from completely random data (Minh et al. 2018). Of note, paraphyly is a well-known artefact when a bifurcating tree is inferred from reticulated species (Sosef 1997; Schmidt-Lebuhn 2012), with concatenation or binning approaches using genomic data being demonstrably vulnerable (Bangs et al. 2018). Thus, we tentatively attribute the observed paraphyly as an artefact of documented hybridization between *G. cypha* and *G. robusta* (Chafin et al. 2019), and the inability of PoMo to model hybridization. Hybridization also potentially drives the lack of monophyly in *G. seminuda,* per TICR and the concatenation tree (Fig. S1).

We also explored the impact of matrix occupancy filters on SVDquartets, and bootstrap support and overall topological consistency declining with increasingly stringent filters (Fig. 3b). This corroborates prior evaluations with regard to the impacts of over-filtering RADseq data (Eaton et al. 2017). In all cases, site-wide concordance was significantly predicted by subtending branch lengths, but not by node depths (Fig. S6). This suggests that site-wise concordance was unbiased in our analyses at either shallower or deeper timescales, but was affected instead by the extent of time separating divergences. Some bioinformatic biases such as ortholog misidentification or lineage-specific locus dropout will disproportionally affect deeper nodes (Eaton 2017). However, we interpret the lack of correlation between node depth and site-wise concordance as an indication that these processes lack substantial bias.

### Biogeographic Hypotheses and Co-divergence

EcoEvolity model selection was not found to be vulnerable to alternative event priors (Fig. S7). The best-fitting model across all priors consistently demonstrated co-divergence of *G. jordani* with the lower basin complex (*G. robusta* x *G. intermedia* and *G. intermedia* x *G. nigra*; Fig. 6). The divergence of *G. elegans* and *G. seminuda* from a theoretical lower basin ancestor pre-dates this putatively rapid radiation, although it is unclear if these estimates were impacted by the aforementioned introgression between *G. seminuda* and *G. elegans*.

**Figure 6:**
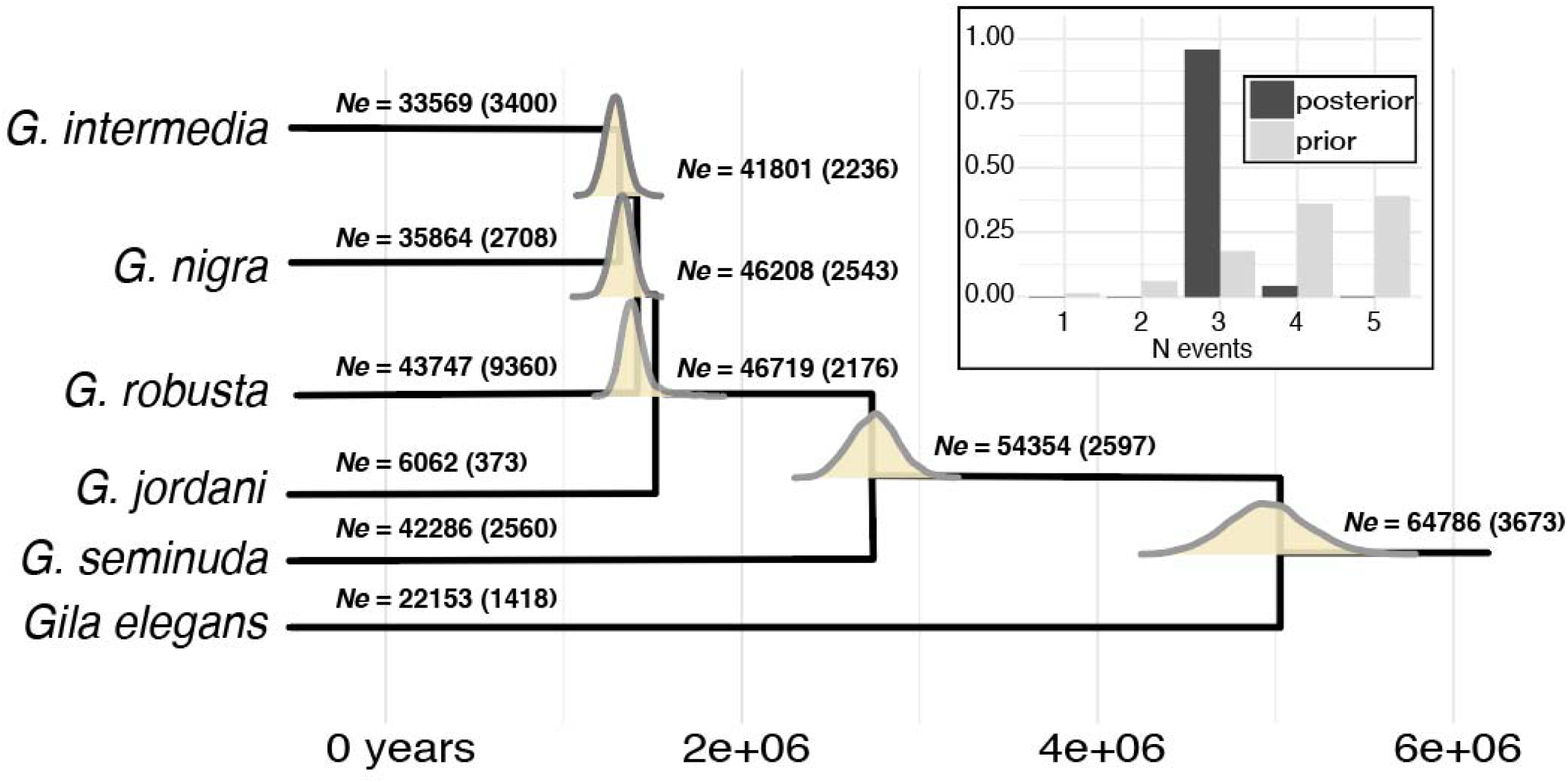
Posterior estimates for divergences times and effective populations sizes (*N*_e_) derived from EcoEvolity and 2,000 randomly sampled full-length ddRAD locus alignments. Branches are annotated with mean (std. dev.) *N*_e_ and posterior probabilities for divergence times are plotted on corresponding nodes. Units are in years, using a static mutation rate of 1.2 e^−08^ substitutions per year. Posterior probabilities for divergence models (insert) suggest the co-divergence of *Gila jordani*, *G. robusta*, *G. nigra*, and *G. intermedia*.

Posterior effective population size (*N*_e_) estimates were large (e.g. >20,000) and consistent with previous estimates (Garrigan et al. 2002). *Gila jordani* was an exception, with a mean posterior *N*_e_=6,062. This discrepancy is not surprising, given the extremely narrow endemism of this species (Tuttle and Scoppettone 1990), and its recent bottleneck (Hardy 1982), although this is still a rather large estimate given the latter. Posterior divergence time estimates suggested a late-Miocene/ early-Pliocene origin of *G. elegans*. Results for *G. seminuda* and the lower basin radiation indicated Pliocene and early Pleistocene divergences, respectively. These results are supported in the fossil record (Uyeno 1960; Uyeno and Miller 1963), although we note paleontological evaluations of *Gila* have been sparse. Thus, we hesitate to interpret these as absolute dates, given our fixed mutation rate for these analyses and an uncertainty regarding the capacity of RADseq methods to yield an unbiased sampling of genome-wide mutation rate variation (e.g. Cariou et al. 2016).

## Discussion

The goal of our study was to determine if extensive geographic and genomic sampling could resolve the taxonomically recalcitrant *G. robusta* complex. We applied diverse phylogenetic models and tests of hybridization and predictions of parameter space within the anomaly zone to diagnose sources of subtree discordance. In so doing, we also tested multiple hypothesized hybrid speciation events. We detected a single reticulation (*G. seminuda*), although other events with a lower component of genomic introgression may have also occurred. We documented rapid co-divergence of lower basin taxa within the anomaly zone and were able to resolve these despite the prevalence of incomplete lineage sorting. This scenario (as outlined below) is consistent with the geomorphology of the region and seemingly represents an adaptive radiation by our study complex, as facilitated by drainage evolution.

### Methodological Artefacts and Conflicting Phylogenetic Hypotheses for Gila

Increased geographic and genomic sampling revealed the presence of diagnosable lineages within the *G. robusta* complex, with both rapid and reticulate divergences influencing inter-locus conflict. Phylogenetic hypotheses for our focal group had previously been generated using allozymes (Dowling and DeMarais 1993), Sanger sequencing (Schwemm 2006; Schönhuth et al. 2014), microsatellites (Dowling et al. 2015), and more recently RADseq (Copus et al. 2018). None could resolve relationships within the lower basin complex. To explain these contrasts, we argue that prior studies suffered from systematic artefacts and ascertainment biases that were overcome, at least in part, by our approach.

Incomplete or biased sampling is a familiar problem for biologists (e.g. Hillis 1998; Schwartz and McKelvey 2009; Ahrens et al. 2016), and we suggest it served as a major stumbling block for delineating the evolutionary history of *Gila*. Unfortunately, insufficient sampling is common in studies of threatened and endangered species, and its repercussions are severe with regard to phylogenetic inference (Hillis 1998). This fact is substantiated by the many examples in which increasingly comprehensive geographic sampling spurred a revision of phylogenetic hypotheses (e.g. Oakey et al. 2004; Linck et al. 2019). Likewise, incomplete sampling of genome-wide topological variation (e.g. Maddison 1997; Degnan and Rosenberg 2009) is an additional source of bias, especially when a very small number of markers are sampled. These issues alone may explain the variation among prior studies. For example, Schwemm (2006) sampled extensively, including nearly all of the sites included in this study, but was only able to examine a handful of genes. Because anomalous gene trees are most probable under a scenario of rapid radiation (as documented herein), the reduced number of loci used by Schwemm (2006) could not recover a consistent species tree. Copus et al. (2016, 2018) examined a dataset containing 6,658 genomic SNP loci (across 1,292 RAD contigs), but only did so across a sparse sample of 19 individuals. A bioinformatic acquisition bias also likely impacted this study, in the form of strict filtering that disproportionately excluded loci with higher mutation rates (Huang and Knowles 2016).

A necessary consideration when validating phylogenetic hypotheses across methods (and datasets) is to gauge compatibility between the underlying evolutionary processes and those actually being modeled (Walker et al. 2018). In this sense, the consideration of statistical support metrics alone can be not only misleading, but also promote false conclusions. For example, bootstrapping is by far the most prevalent method of evaluating support in phylogenetic datasets (Felsenstein 1985). While bootstrap concordances may be appropriate for moderately-sized sequence alignments (e.g. Efron et al. 1996), they can be meaningless when applied to sufficiently large datasets (Gadagkar et al. 2005; Kumar et al. 2012). This is apparent in the high bootstrap support displayed for anomalous relationships in our analysis (Fig. S1). Phylogenetic signal also varies among loci, such that in many instances relatively few loci drive contentious relationships (Shen et al. 2017). Likewise, not all methods are equal with respect to their simplifying assumptions. Given this, we deem it imperative to consider the biases and imperfections in both our data, and the models we apply.

### Complex Evolution and Biogeography of the Colorado River

The taxonomic instability in *Gila* is not uncommon for fishes of western North America, where confusing patterns of diversity were generated by tectonism and vulcanism (Minckley et al. 1986; Spencer et al. 2008). This issue is particularly emphasized when viewed through the lens of modern drainage connections (Douglas et al. 1999). Historic patterns of drainage isolation and intermittent fluvial connectivity not only support our genomic conclusions but also summarize the paleohistory of the Colorado River over temporal and spatial scales.

The earliest record of fossil *Gila* from the ancestral Colorado River is mid-Miocene (Uyeno and Miller 1963), with subsequent Pliocene fossils representing typical ‘big river’ morphologies now associated with *G. elegans*, *G. cypha*, and *G. robusta* (Uyeno and Miller 1965). The modern Grand Canyon region lacked any fluvial connection at the Miocene-Pliocene transition, due largely to regional tectonic uplifts that subsequently diverted the Colorado River (Spencer et al. 2001; House et al. 2005). Flows initiated in early Pliocene (c.a. 4.9 mya; Sarna-Wojcicki et al. 2011), and subsequently formed a chain of downstream lakes associated with the Bouse Formation (Lucchitta 1972; Spencer and Patchett 2002). Evidence suggests ‘spillover’ by a successive string of Bouse Basin paleolakes was episodic, and culminated in mid-Pliocene (House et al. 2008), with an eventual marine connection via the Salton Trough to the Gulf of California (Dorsey et al. 2007). Prior to this, the Gila River also drained into the Gulf (Eberly and Stanley 1978), and sedimentary evidence indicated that it was isolated from the Colorado until at least mid-Pliocene by a northward extension of the Gulf (Helenes and Carreno 2014). This geomorphology is reflected in a broader phylogeographic pattern that underscores marked differences between resident fish communities in the upper and lower basins (Hubbs and Miller 1948).

Intra-basin diversification also occurred as an addendum to hydrologic evolution. Although the course of the pluvial White River is now generally dry, it may have been a Pliocene-early Pleistocene tributary of a paleolake system when the proto-Colorado River first extended into the modern-day lower basin (Dickinson 2013). This may represent an initial colonization opportunity for upper basin fishes, an hypothesis that coincidentally aligns well with our rudimentary age estimate for Virgin River chub, *G. seminuda* (Fig. 6). This early isolation, as well as the continued contrast between the spring-fed habitats therein, and the high flows of the ancestral Colorado River, provide an explanation for the unique assemblage of *Gila* and other fishes therein (Hubbs and Miller 1948). Thus, as has been shown to be the case in other taxa, the biogeographic context (here in the form of drainage evolution) is likely an important factor in the apparent reticulate evolution in *Gila* (Burbrink and Gehara 2018).

Phylogenetic signatures of the anomaly zone (Fig. 5) coupled with co-divergence modeling (Fig. 6) suggest the diversification of lower basin *Gila* occurred rapidly post-colonization. Late Pliocene integration of the two basins provided an opportunity for dispersal into the lower basin tributaries. The Plio-Pleistocene climate of the region was quite different, with a relatively mesic Pliocene as precursor to a protracted monsoonal period extending through early Pleistocene (Thompson 1991; Smith et al. 1993). The latter, in turn, may have resulted in relatively unstable drainage connections (Huckleberry 1996). The potential for climate-driven instability, and the complex history of intra-drainage integration of Gila River tributaries during the Plio-Pleistocene (Dickinson 2015), lends support to the ‘cyclical-vicariance’ model proposed by Douglas et al. (1999). These periods of isolation may have promoted an accumulation of ecological divergences that persisted post-contact, and were sufficient to maintain species boundaries despite contemporary sympatric distributions and weak morphological differentiation. This hypothesis is also supported by the non-random mating found among *G. robusta* and *G. nigra,* despite anthropogenically-induced contact (Marsh et al. 2017).

### Management Implications

A request by the Arizona Game and Fish Department to review the taxonomy of the *Gila robusta* complex prompted the American Fisheries Society (AFS) and the American Society of Ichthyology and Herpetology (ASIH) to recommend the synonymization of *G. intermedia* and *G. nigra* with *G. robusta*, owing in part to their morphological ambiguity and an imprecise taxonomic key (Carter et al. 2018). Given this, a proposal to extend protection to lower basin *G. robusta* and *G. nigra* at the federal level was subsequently withdrawn (USFWS 2017; Fig. 1). As was the case prior to this withdrawal, *G. intermedia* alone is classified as endangered (USFWS 2005) under the Endangered Species Act (ESA 1973; 16 U.S.C. § 1531 et seq).

This study provides a much needed resolution to this debate by defining several aspects: First, our study reinforced the recognition of *G. robusta* as demonstrably polyphyletic, with two discrete, allopatric clades corresponding to the upper and lower basins of the Colorado River (Dowling and DeMarais 1993; Schönhuth et al. 2014). These data, together with the geomorphic history of the region that promoted endemic fish diversification (as above), clearly reject ‘*G. robusta*’ as a descriptor of contemporary diversity. This underscores a major discrepancy in the taxonomic recommendations for the lower basin complex (Page et al. 2016). Given that the type locality of *G. robusta* is in the upper basin (i.e., the Little Colorado River), we note a pressing need either to determine taxonomic precedence for the lower basin ‘*G. robusta,*’ or to provide a novel designation. The potential resurrection of a synonym is a possibility, necessitating detailed examinations of the type specimens prior to a formal recommendation. This may be appropriately adjudicated by the AFS-ASIH Names of Fishes Committee, as a follow-up to their earlier involvement.

The situation with *G. intermedia* and *G. nigra* is slightly more ambiguous. The short internodes and anomaly zone divergences identified herein explain previous patterns found in population-level studies, with elevated among-population divergence but scant signal uniting species (Dowling et al. 2015). We also unequivocally rejected the previous hypothesis of hybrid speciation for *G. nigra* (Minckley and DeMarais 2000; Dowling et al. 2015).

Rather, intermediacy in the body shape of *G. nigra* reflects differences accumulated during historic isolation (Douglas et al. 1999) and/ or the retention of an adaptive ecomorphology (Douglas and Matthews 1992). These hypotheses warrant further exploration, with provisional results employed in future management decisions (Forest et al. 2015). With regards to taxonomy, we confidently recommend that *G. intermedia* be resurrected, and that additional studies be implemented to dissect the potential distinctiveness of *G. nigra*. For management purposes, we echo a conservative, population-centric approach (previously argued for by Dowling et al. 2015; Marsh et al. 2017).

Three primary components of a ‘Darwinian shortfall’ in biodiversity conservation are recognized (Diniz-Filho et al. 2013): (i) The lack of comprehensive phylogenies; (ii) Uncertain branch lengths and divergence times; and (iii) insufficient models linking phylogenies with ecological and life-history traits. Taxonomic uncertainty in *Gila* is severely impacted by the first two of these, with taxonomic resolution prevented by the comingling of sparse phylogenetic coverage with temporal uncertainty. We must now address the relationships between ecology, life history, and phylogeny in *Gila,* so as to understand the manner by which phylogenetic groupings (identified herein) are appropriate as a surrogate for adaptive/ functional diversity. For example: To what degree are *Gila* in the lower basin ecologically non-exchangeable? How do they vary in their respective life histories? Is reproductive segregation maintained in sympatry (as in Marsh et al. 2018), and if so, by what mechanism?

## Conclusion

The intractable phylogenetic relationships in *Gila* were resolved herein through improved spatial and genomic sampling. Our data, coupled with polymorphism-aware methods and contemporary approaches that infer trees, yielded a revised taxonomic hypothesis for *Gila* in the lower Colorado River basin. The geomorphic history of the Colorado River explains many anomalous patterns seen in this and previous studies, wherein opportunities for contact and colonization were driven by the tectonism characteristic for the region. The signal of rapid diversification is quite clear in our data, as interpreted from patterns inherent to phylogenetic discord. We emphasize that discordance in this sense does not necessarily represent measurement error or uncertainty. Instead, it is an intrinsic component of phylogenetic variance that is not only expected within genomes (Maddison 1997), but also a necessary component from which to build hypotheses regarding the underlying evolutionary process (Hahn and Nakhleh 2016). Ignoring this variance in pursuit of a ‘resolved phylogeny’ can lead to incorrect inferences driven by systematic error. Similarly, insufficient spatial or genomic sampling may also promote a false confidence in anomalous relationships, particularly when character sampling is particularly dense, whereas taxon sampling is sparse.

We reiterate that phylogenetic hypotheses, by their very nature, cannot exhaustively capture the underlying evolutionary process. One approach is to categorize phylogenetic (and ‘non-phylogenetic’) signals in those regions of the tree that are refractive to certain models (as done herein). We also acknowledge that attempting to reconstruct the past using contemporary observations is a battle against uncertainty and bias, with the revisions of phylogenetic/ taxonomic hypotheses expected as additional data are accrued. As such, we urge empiricists that engage in taxonomic controversies (such as this one) to interrogate their results for transparency. The task of sorting through conflicting recommendations invariably falls to natural resource managers, with unreported biases (be they methodological or geopolitical) only confounding those efforts.

## Acknowledgements

This research was conducted in partial fulfillment by TKC of the Ph.D. degree in Biological Sciences at the University of Arkansas, as enabled by a Distinguished Doctoral Fellowship (DDF) award. Numerous agencies and organizations contributed field expertise, sampling, permit authorization, funding, and/or valuable comments: Arizona Game and Fish Department, Colorado Parks and Wildlife, Jicarilla Apache Game and Fish, Nevada Department of Wildlife, New Mexico Department of Game and Fish, United States Fish and Wildlife Service, United States Forest Service, National Park Service, United States Bureau of Reclamation, Utah Department of Natural Resources, Utah Division of Wildlife, and the Wyoming Game and Fish Department. Particular thanks are extended to: J. Alves, M. Anderson, R. Anderson, P. Badame, K. Bestgen, M. Breen, M. Brouder, K. Breidinger, S. Bryan, P. Cavalli, B. DeMarais, T. Dowling, R. Fridell, K. Gelwicks, K. Guadalupe, J. Harter, B. Harvey, K. Hilwig, M. Hudson, D. Keller, J. Logan, C. McAda, S. Meiser, C. Minckley, T. Modde, K. Morgan, G. Munsen, F. Pfeifer, W. Radke, A. Rehm, S. Ross, M. Smith, R. Stevenson, R. Timmons, S. Tolentino, P. Unmack, A. Varelas, D. Weedman, K. Wilson, and E. Woodhouse (with apologies to anyone inadvertently overlooked). We also received tissues from the Bell Museum at the University of Minnesota (JJPE 06-15, 06-16, 06-20, 06-24), Los Angeles County Museum of Natural History (LACM 555990-1, 57271-1), Monte L. Bean Life Science Museum at Brigham Young University (BYU 57580-4, 68470-4, 138751-2 61643-8), Museum of Southwestern Biology at the Univeristy of New Mexico (MSB 44631, 81351, 81493), and Oregon State University Ichthyology Collection (OSU 15444, 17901), for which we thank R. Feeney, D. Shiozawa, B. Sidlauskas, A. Simons, A. Snyder and the original collectors of accessioned materials. Additional sampling was completed by MED and MRD under permits provided by: Arizona Game and Fish Department, The Hualapai Tribe, The Navajo Nation, Nevada Department of Wildlife, U.S. National Park Service, and Wyoming Department of Game and Fish. Federal permit TE042961-0 (1434-HQ-97RU-01552 RWO #66) was provided by the U.S. Game and Fish Service. Sampling procedures were approved under Arizona State University Animal Care and Use Committee (ASU IACUC) permit 98–456R and Colorado State University Animal Care and Use Committee (CSU IACUC) permit 01–036A-01. Travel in the Grand Canyon was conducted under auspices of Grand Canyon National Park river use permits. We are also indebted to students, postdoctorals, and faculty who have promoted our research: A. Alverson, W. Anthonysamy, M. Davis, L. James, J. Pummill, A. Tucker. Funding was provided by several generous endowments from the University of Arkansas: The Bruker Professorship in Life Sciences (MRD), and the Twenty-First Century Chair in Global Change Biology (MED). This work was also in part facilitated by funding provided by the Arizona Game and Fish Heritage Fund, Colorado Parks and Wildlife, and U.S. Geological Survey Species at Risk Program. Additional analytical resources were provided by the Arkansas Economic Development Commission (Arkansas Settlement Proceeds Act of 2000) and the Arkansas High Performance Computing Center (AHPCC), and from an NSF-XSEDE Research Allocation (TG-BIO160065) to access the Jetstream cloud. The use of trade, product, industry or firm names is for informative purposes only and does not constitute an endorsement by the U.S. Government or the U.S. Fish and Wildlife Service. Links to non-Service Web sites do not imply any official U.S. Fish and Wildlife Service endorsement of the opinions or ideas expressed therein or guarantee the validity of the information provided. The findings, conclusions, and opinions expressed in this article represent those of the authors, and do not necessarily represent the views of the U.S. Fish & Wildlife Service.

## Competing Interests

The authors declare no conflict of interest.

## Data Accessibility

Pending acceptance, all relevant (i.e. assembled and filtered) datasets will be archived on Dryad. Codes and custom scripts developed in support of this work are available as open-source under the GNU Public License via GitHub: github.com/tkchafin

- Sampling locality metadata and alignments: Dryad (pending)
- Codes and scripts: github.com/tkchafin (and as cited in-text)

## Author Contributions

All authors constributed to research conceptualization and design. MED and MRD contributed sampling and coordination of agency efforts. TKC performed molecular work and analyses. TKC, SMM, BTM, and MRB wrote relevant codes and contributed analyses. All authors contributed to writing the manuscript and approve of the final submission.

## Supplementary Materials

**Table S1:**
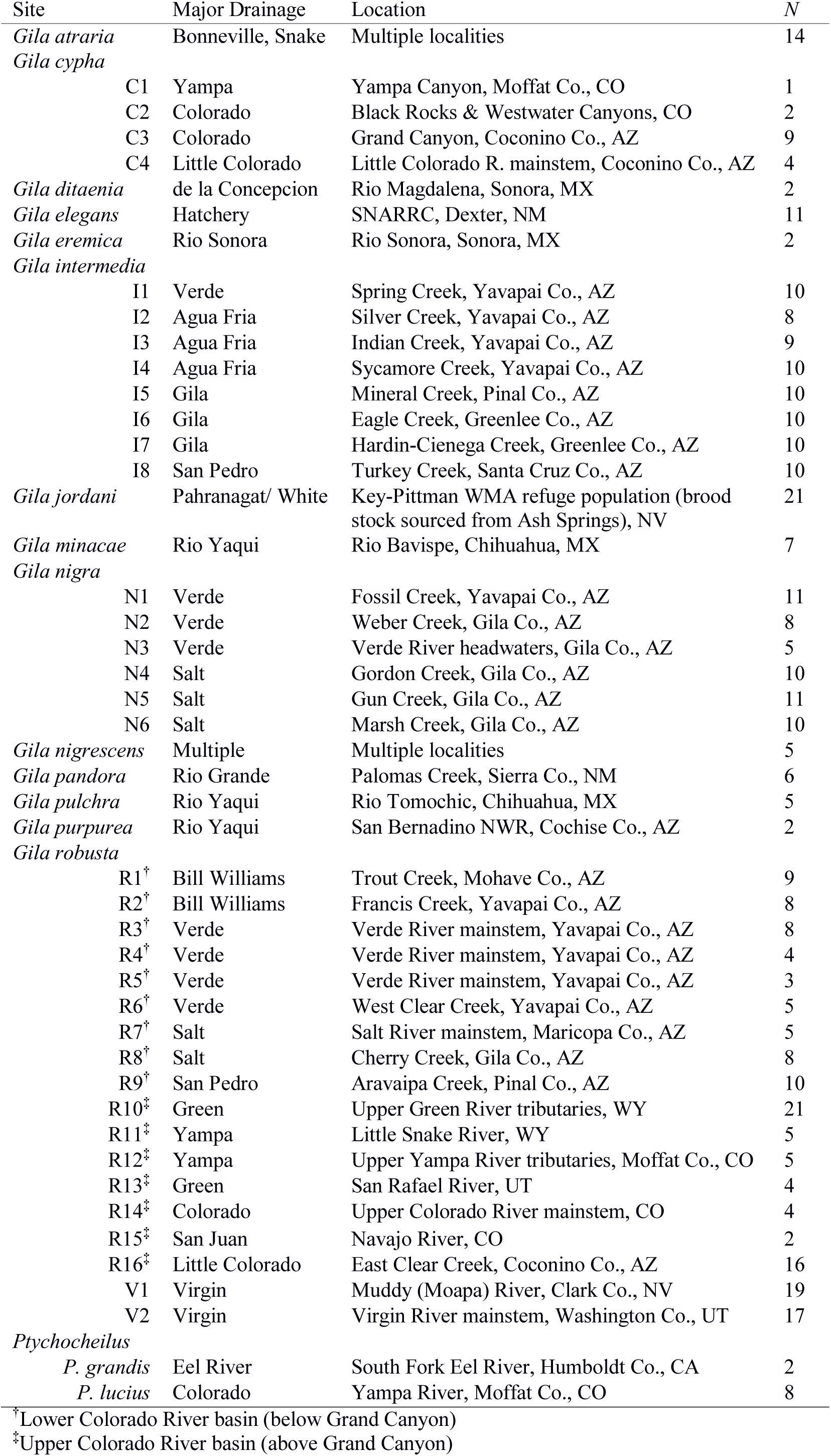
Sampling locations and drainages for *N*=386 *Gila* individuals and outgroups.

**Figure S1:**
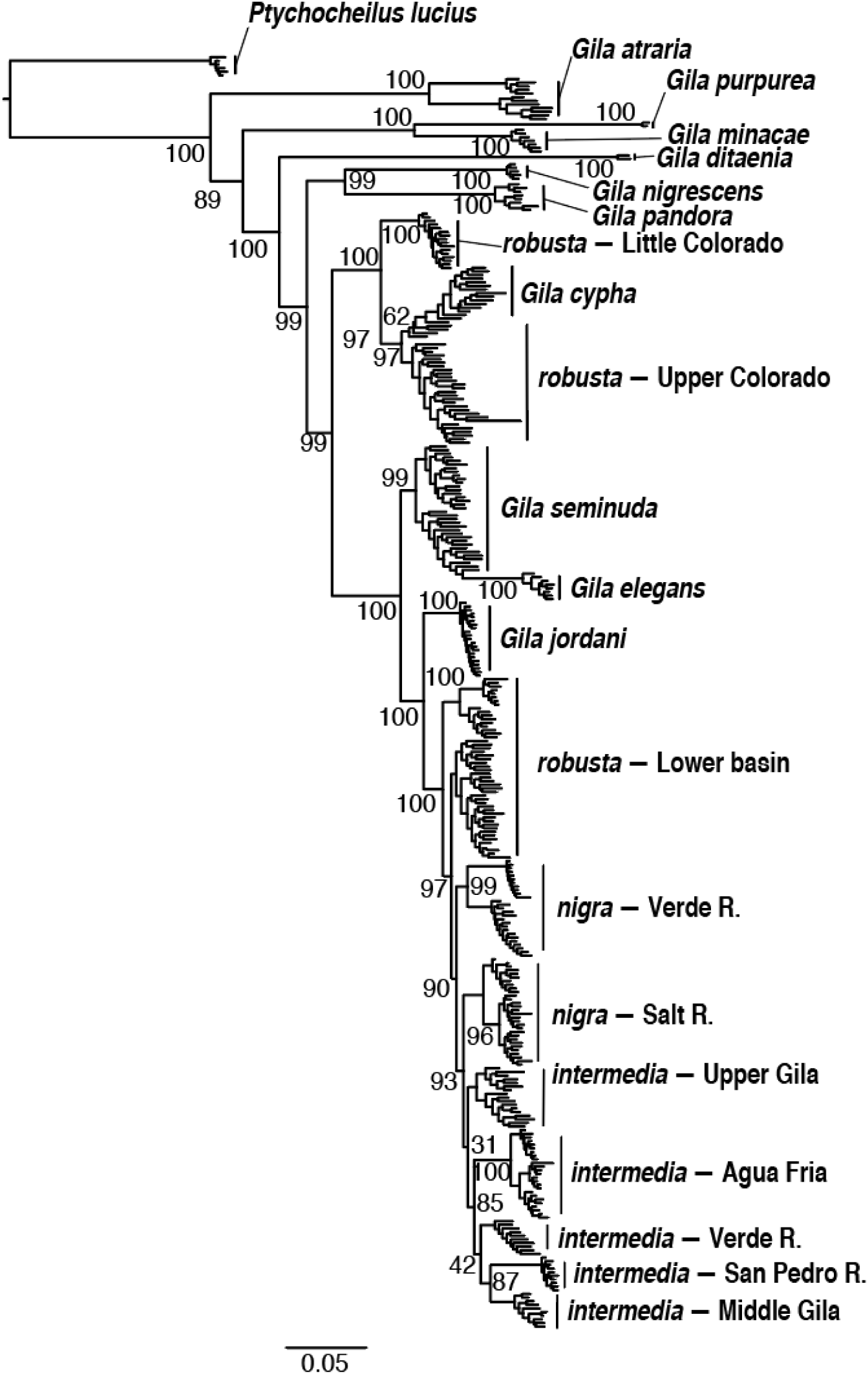
Phylogram showing results from an unconstrained search using 21,717 concatenated SNPs in Iq-Tree. Focal nodes are annotated with bootstrap support (values for shallow nodes omitted for clarity). For specific locality information, refer to Table S1.

**Figure S2:**
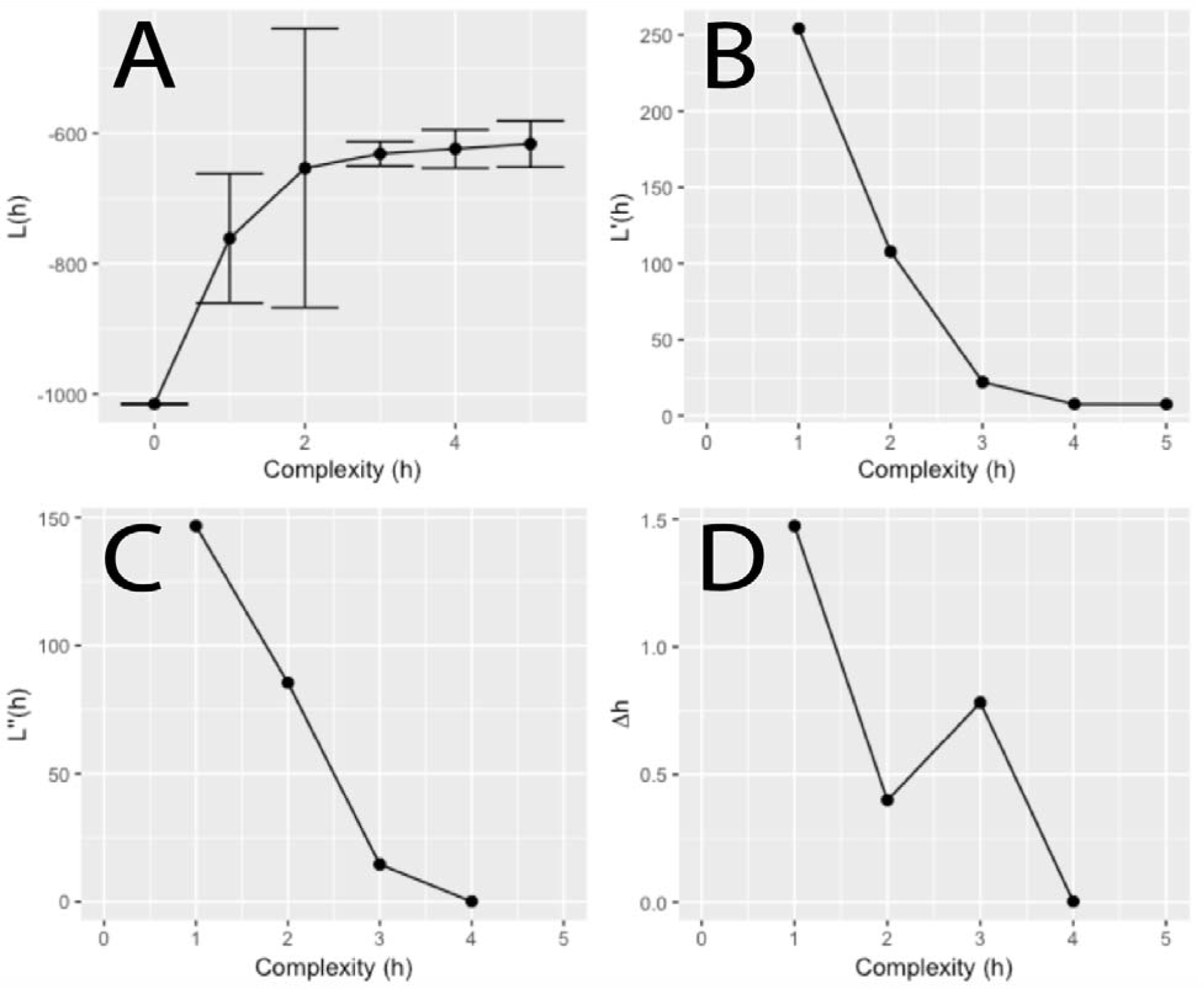
Model selection results for SNaQ/ PhyloNetworks; *h*=maximum number of hybrid edges allowed per model; (A) L(*h*) = −log likelihood for the best network of *N*=48 replicate runs per value of *h*; (B) L’(*h*) = 1^st^ order change in L(*h*) = L(*h*) – L(*h*-1); (C) L’’(*h*) = 2^nd^ order change in L(*h*) = L’(*h*+1) – L’(*h*); and (D) Δ*h* = L’’(*h*) / s(*h*) where s(*h*) is the standard deviation in L(*h*) among replicates.

**Figure S3:**
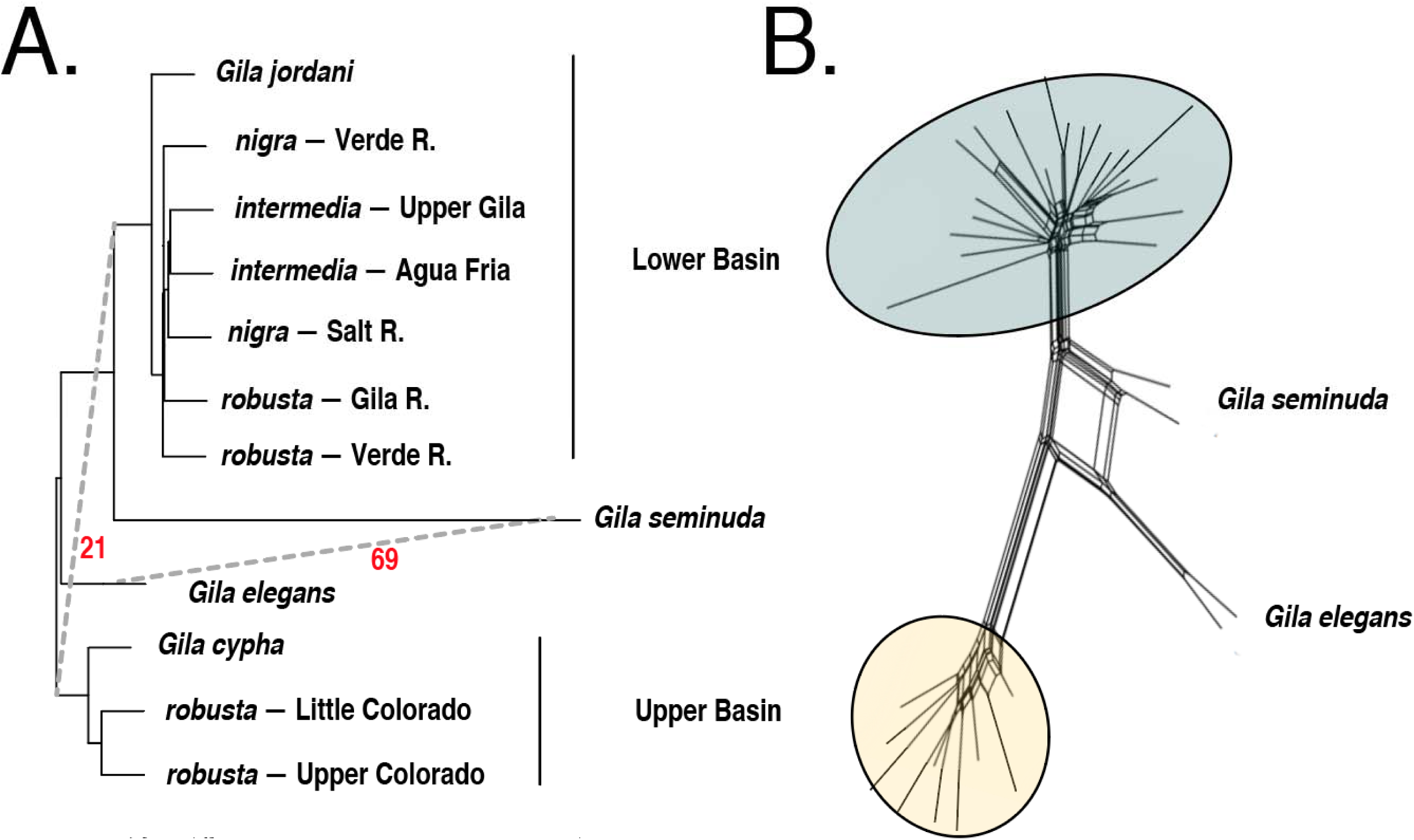
Reticulate networks inferred using (A) 100 bootstrap pseudoreplicates in the SNaQ algorithm and (B) the distance-based NeighborNet algorithm in SplitsTree4.

**Figure S4:**
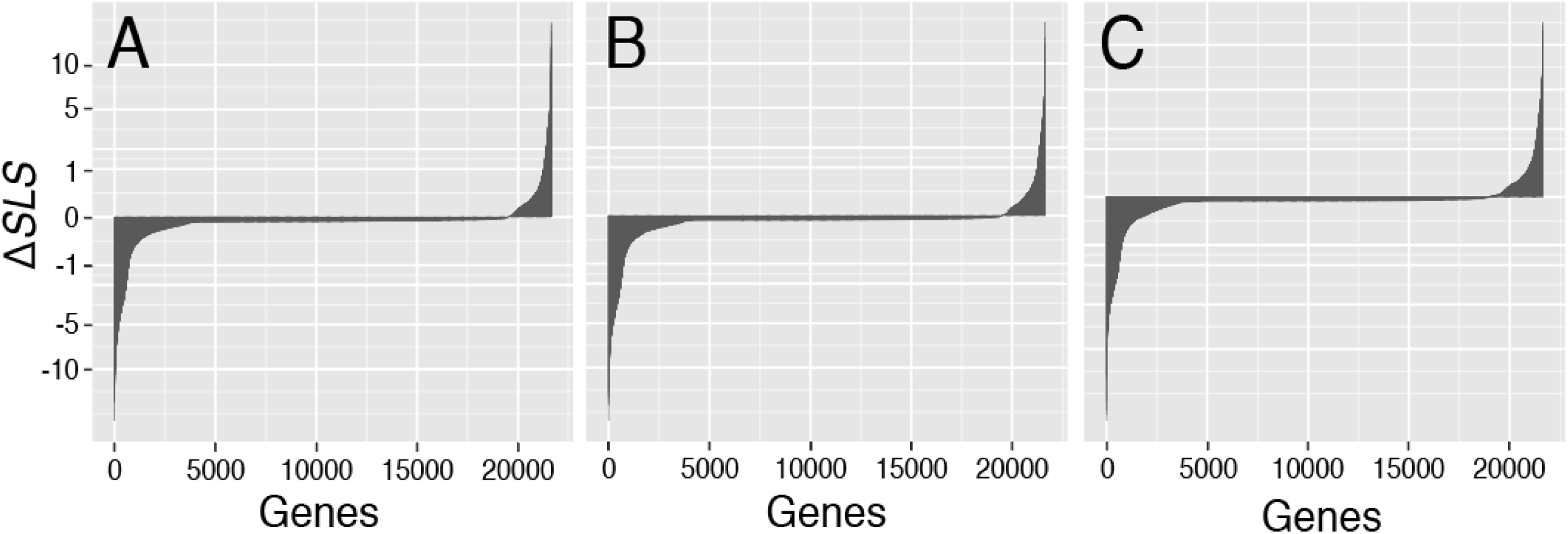
Site-wise log-likelihood differences (Δ*SLS*) for (A) SVDQuartets, (B) PoMo, and (C) TICR topologies as compared to an unconstrained concatenated tree. Δ*SLS* values are transformed as signed square-roots, with positive values indicating increased site-likelihood under the constrained model, and negative values having increased likelihood under the unconstrained concatenated model.

**Figure S5:**
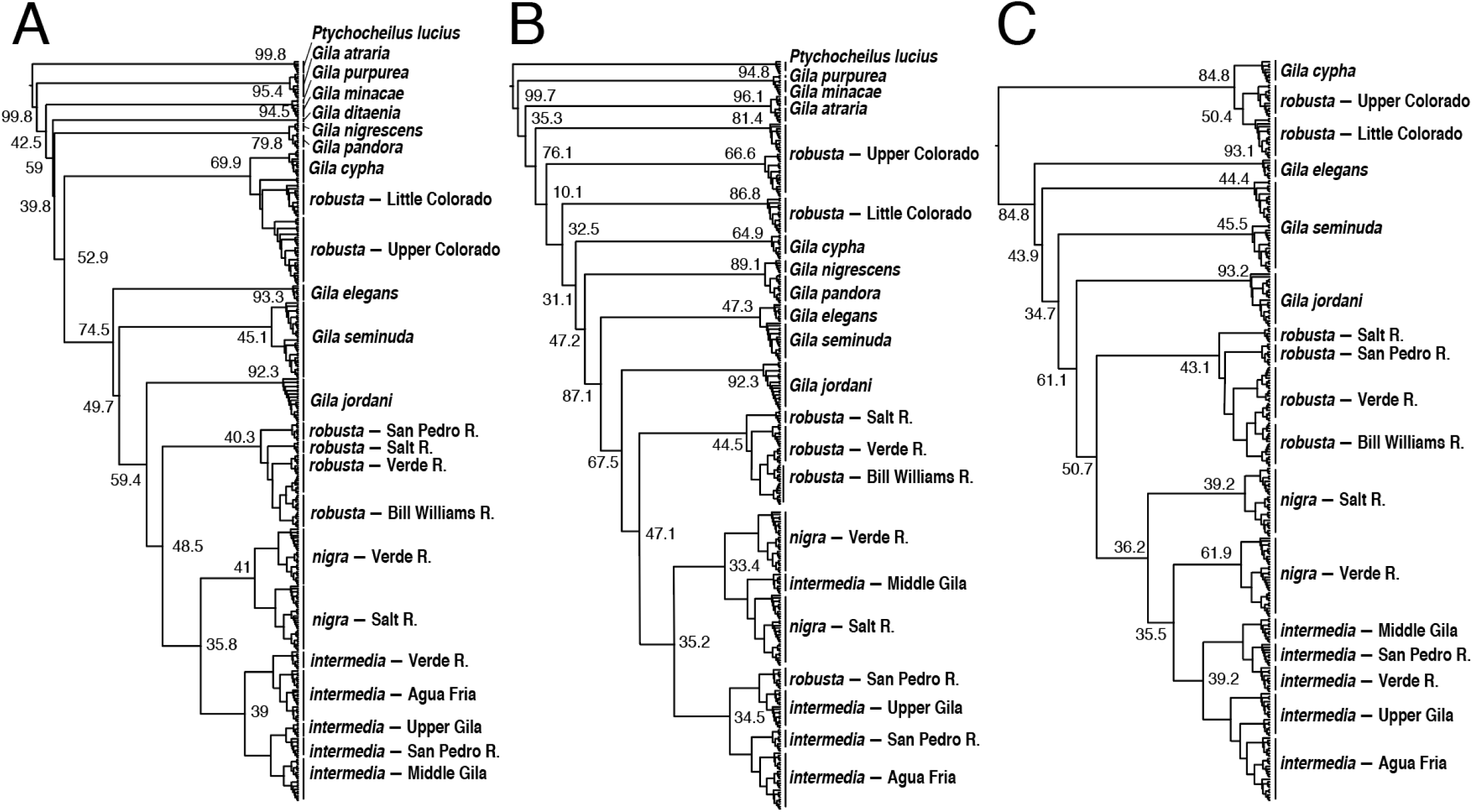
Site-wise concordance factors (s*CF*) for lineage trees produced in Iq-Tree under topological constraints for the (A) SVDQuartets, (B) PoMo, and (C) TICR results. For details, see Methods.

**Figure S6:**
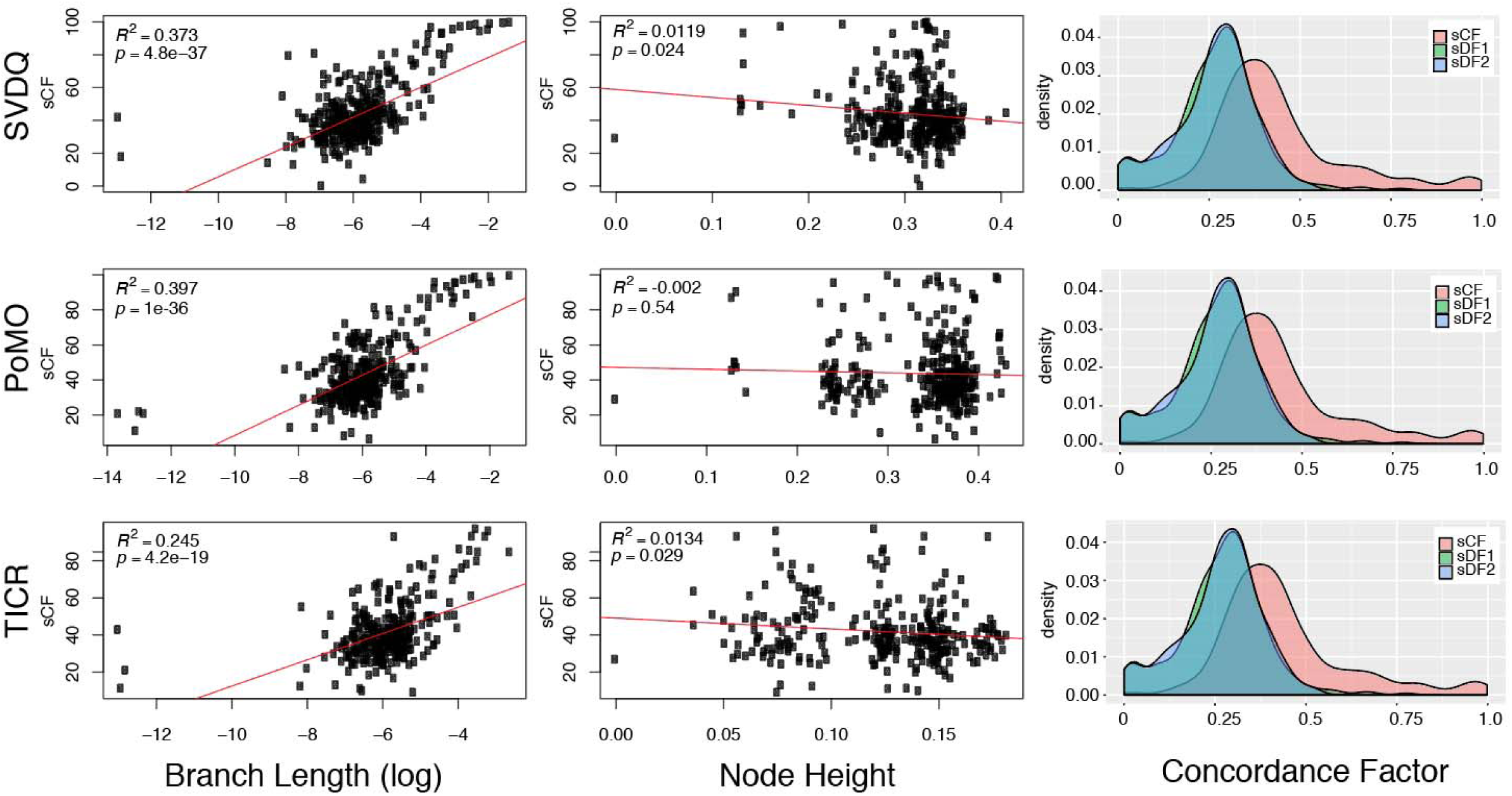
Characterization of site-wise concordance (*s*CF) factors for SVDQuartets, PoMo, and TICR phylogenies. Panels show (left to right): Linear regression of subtending branch lengths (log-transformed) with *s*CF; node height (cumulative branch lengths from root to focal node); and densities of *s*CF across nodes as compared to the discordance factors for the two conflicting quartet resolutions (*s*DF1 and *s*DF2).

**Figure S7:**
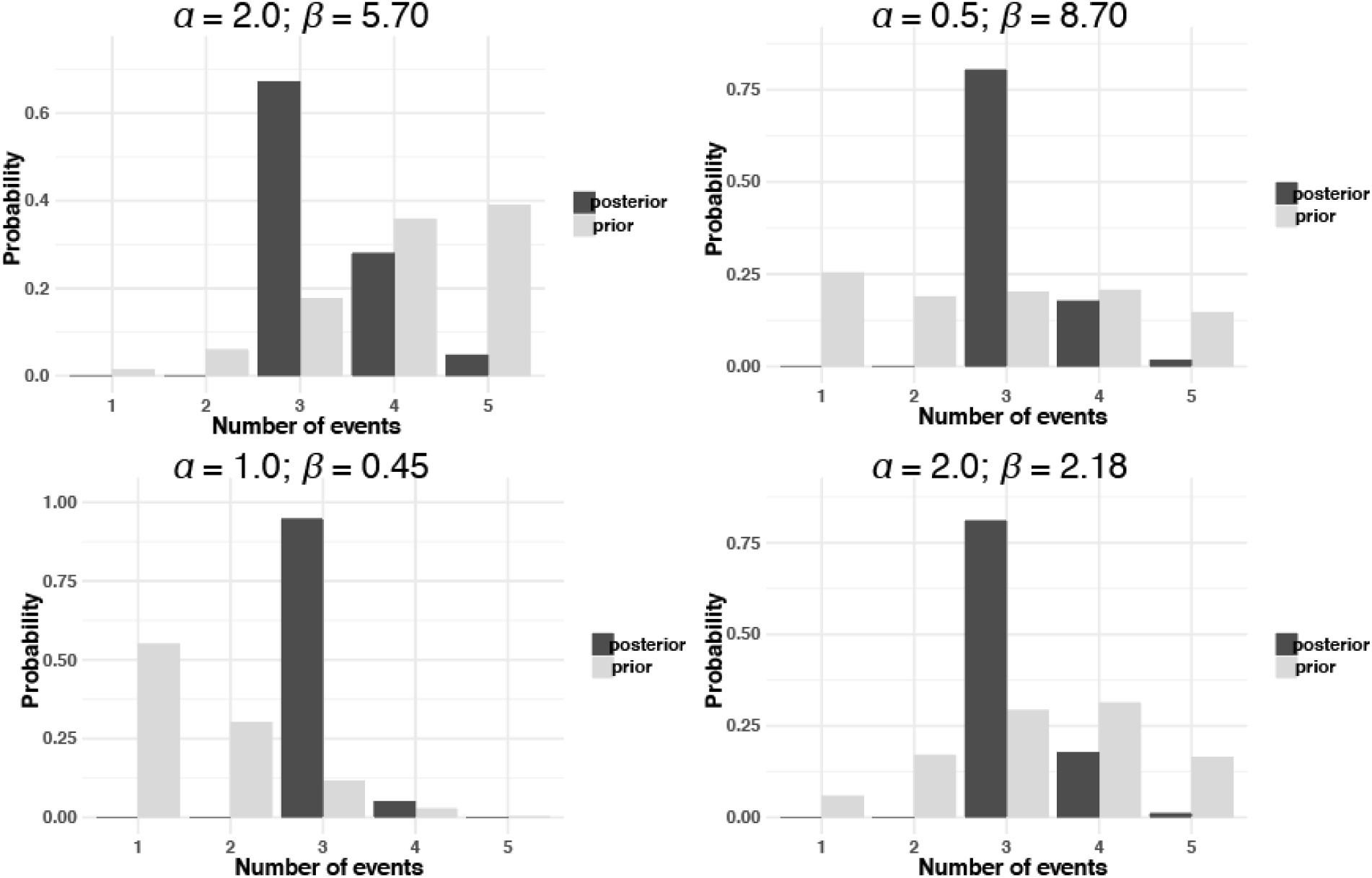
Prior and posterior probabilities for number of independent divergence events in EcoEvolity co-divergence models for *Gila*. Parameters across all runs were identical, except for the shape (*α*) and scale (*β*) of the gamma-distributed prior on the Dirichlet process concentration.

## Notes

### Competing Interest Statement

The authors have declared no competing interest.

### Summary of Updates

Manuscript was rejected with major revision from the journal Systematic Biology; this version addresses those requested revisions.

